# Oxidative-phosphorylation genes from the mitochondrial and nuclear genomes co-express in all human tissues except for the ancient brain

**DOI:** 10.1101/136457

**Authors:** Gilad Barshad, Amit Blumberg, Dan Mishmar

**Affiliations:** Department of Life Sciences, Ben-Gurion University of the Negev, Beer Sheva 8410501, Israel

**Keywords:** gene expression, mitochondria, mito-nuclear, oxidative phosphorylation, RNA-seq, tissue specificity

## Abstract

In humans, oxidative phosphorylation (OXPHOS), the cellular energy producer, harbors ∼90 nuclear DNA (nDNA)- and mitochondrial DNA (mtDNA)-encoded subunits. Although nDNA- and mtDNA-encoded OXPHOS proteins physically interact, their transcriptional regulation profoundly diverges, thus questioning their co-regulation. To address mtDNA-nDNA gene co-expression, we analyzed ∼8,500 RNA-seq Gene-Tissue-Expression (GTEx) experiments encompassing 48 human tissues. We found overall positive cross-tissue mtDNA-nDNA OXPHOS gene co-expression. Nevertheless, alternatively-spliced variants, as well as certain OXPHOS genes, did not converge into the main OXPHOS gene cluster, suggesting tissue-specific flavor of OXPHOS gene expression. Finally, unlike non-brain body sites, and neocortex and cerebellum (‘mammalian’ brain), negative mito-nuclear expression correlation was found in the hypothalamus, basal ganglia and amygdala (‘ancient brain’). Analyses of co-expression, DNase-seq and ChIP-seq experiments identified candidate RNA-binding genes and CEBPb as best explaining this phenomenon. We suggest that evolutionary convergence of the ‘mammalian’ brain into positive mtDNA-nDNA OXPHOS co-expression reflects adjustment to novel bioenergetics needs.

## Introduction

Energy production via mitochondrial oxidative phosphorylation (OXPHOS) is the major source of energy in most human tissues. OXPHOS employs five multi-subunit protein complexes harboring more than 80 nuclear DNA (nDNA)-encoded proteins, as well as 13 mitochondrial DNA (mtDNA)-encoded subunits ^1^. As many OXPHOS genes were transferred from the ancestral mitochondrial genome to the host nuclear genome during the course of evolution, these genes are currently located throughout the human karyotype and are individually regulated ^2^. In contrast, human mtDNA-encoded genes have retained their prokaryotic ancestral co-transcription strategy in heavy- and light-strand polycistrons ^3^. These two polycistrons dramatically differ in their coding content, with the first harboring 12 of the 13 OXPHOS subunits, and the remaining protein, the ND6 subunit of OXPHOS complex I, NADH ubiquinone oxidoreductase, being encoded by the light strand. Such prokaryotic-like expression is thought to be regulated mainly by a dedicated mitochondrially targeted RNA polymerase (POLRMT) and two transcription factors (TFAM and mtTF2B) ^4^. Although several known regulators of nuclear gene transcription were recently shown to be imported into the mitochondria and regulate mtDNA transcription ^5–8^, the dispersal of OXPHOS genes throughout the human genome, in addition to their division between the nuclear and cytoplasmic genomes, apparently interferes with transcriptional co-regulation.

Are OXPHOS gene transcripts co-regulated? It was previously argued that genes encoding factors that belong to the same pathway tend to be co-regulated at the transcriptional level ^9,10^. This hypothesis has been tested for nDNA OXPHOS genes at the level of gene expression pattern using Affymetrix chips ^11^. In that study, the authors argued for the tendency of genes encoding OXPHOS subunits to co-express at the mRNA level. The authors further argued that such co-expression tended to be subdivided according to the five OXPHOS protein complexes, i.e., complex I, succinate dehydrogenase (complex II), cytochrome bc1 (complex III), cytochrome oxidase (complex IV) and ATP synthase (complex V). Nevertheless, although the gene expression chip profiles considered represented multiple human tissues (N=79), they were derived from only few individuals and, therefore, cannot provide real insight into OXPHOS gene co-expression in the different tissues. Additionally, as these experiments did not employ mtDNA probes and did not consider tissue-specific splice variants, gaps in our understanding of OXPHOS gene regulation across and within tissues remain. Other analysis that included mtDNA genes was limited to only a few genes ^12^. Recent analysis of RNA-seq data from the Cancer Genome Atlas (TCGA) sample collection enabled Reznik et al. (2017) to assess the expression patterns of mtDNA-encoded OXPHOS subunits and corresponding nDNA-encoded OXPHOS genes in a variety of cancer types and in normal adjacent samples ^13^. Although co-expression of OXPHOS subunits encoded by the two genomes was identified in several cancer types, one cannot easily extend these observations to describe the pattern of OXPHOS gene expression across normal healthy tissues.

Despite the fundamental role played by mitochondrial bioenergetics in eukaryotic metabolism, tissue differences were reported in mitochondrial number, morphology and network interactions ^14^. Such variation, along with tissue diversity in terms of bioenergetics requirements, interferes with any generalization concerning OXPHOS gene co-regulation to all tissues. Indeed, variation in OXPHOS function across tissues was demonstrated in several model organisms ^15^. Furthermore, the identification of tissue-specific subunit paralogs in cytochrome c-oxidase (complex IV) ^16,17^, as well as recent identification of tissue-specific splice variants in the complex I subunit NDUFV3 ^18–20^, provided the first clues of possible tissue variation, and even tissue-specificity, in OXPHOS subunit composition. Although these pieces of evidence support variation in OXPHOS gene expression across tissues, the extent of this phenomenon has yet to be assessed.

Here, we utilized the human RNA-seq gene expression GTEx database ^21,22^, containing experimental information collected from multiple body sites from over 500 donors, to assess co-expression patterns of the entire set of nDNA- and mtDNA-encoded OXPHOS genes across a variety of body sites (N=48). Overall tissue expression patterns were correlated between the two genomes, and candidate transcriptional and post-transcriptional regulatory factors that could best explain the observed co-expression were identified. Unexpectedly, negative mito-nuclear gene expression correlation in ‘ancient’ brain tissues was found. These results underline a profoundly different mode of mitochondrial gene expression regulation in brain tissues associated with the ancient ‘reptile’ brain (i.e, hypothalamus, basal ganglia, amygdala and upper spinal cord), in stark difference to all other body sites, as well as to brain regions that were specifically expanded during mammalian evolution (i.e., cortex, frontal cortex, cerebellar hemispheres, and cerebellum) ^23,24^. Finally, our analysis unearthed a group of structural OXPHOS genes and splice variants that co-express in a tissue-dependent manner, supporting the existence of tissue-specific expression of certain OXPHOS genes.

## Results

### nDNA OXPHOS structural genes are generally co-expressed

Since proteins that participate in the same biochemical activity tend to co-express at the RNA level ^9,10^, we hypothesized that mtDNA and nDNA-encoded OXPHOS genes (Supplementary Table 1) would likely be co-expressed. To test this hypothesis, we assessed correlation between OXPHOS gene expression across multiple human body sites using the publically available GTEx RNA-seq expression portal. For the sake of consistency, and to avoid sample size-related effects, we only considered those body sites for which expression data from at least 50 samples from unrelated donors was available (N=48 tissues), and extracted expression data for OXPHOS genes, which were then normalized for age, gender and cause of death (Supplementary Fig. 1). Next, we calculated expression pattern correlations (Spearman’s rank correlation efficient (r_s_)) for all possible OXPHOS gene pairs across the selected 48 human body sites. A thousand replicates of this procedure enabled calculation of the median Spearman’s rank correlation coefficient. When a significant correlation (p<0.05) was achieved in more than 950 (95%) of the iterations, we considered the correlation to be significant. These calculations generated clusters of co-expressed OXPHOS genes (Supplementary Table 2), here presented as a heat map (Fig. 1). Firstly, such analysis revealed high similarity in the expression pattern of all nDNA-encoded OXPHOS genes, excluding eight genes with known paralogs (i.e., COX6A1, COX6A2, COX4I2, COX7A1, COX8C, COX7B2, COX6B2 and ATP5L2). This further supported the co-expression of nDNA-encoded OXPHOS genes, as previously suggested ^11^.

**Figure 1:**
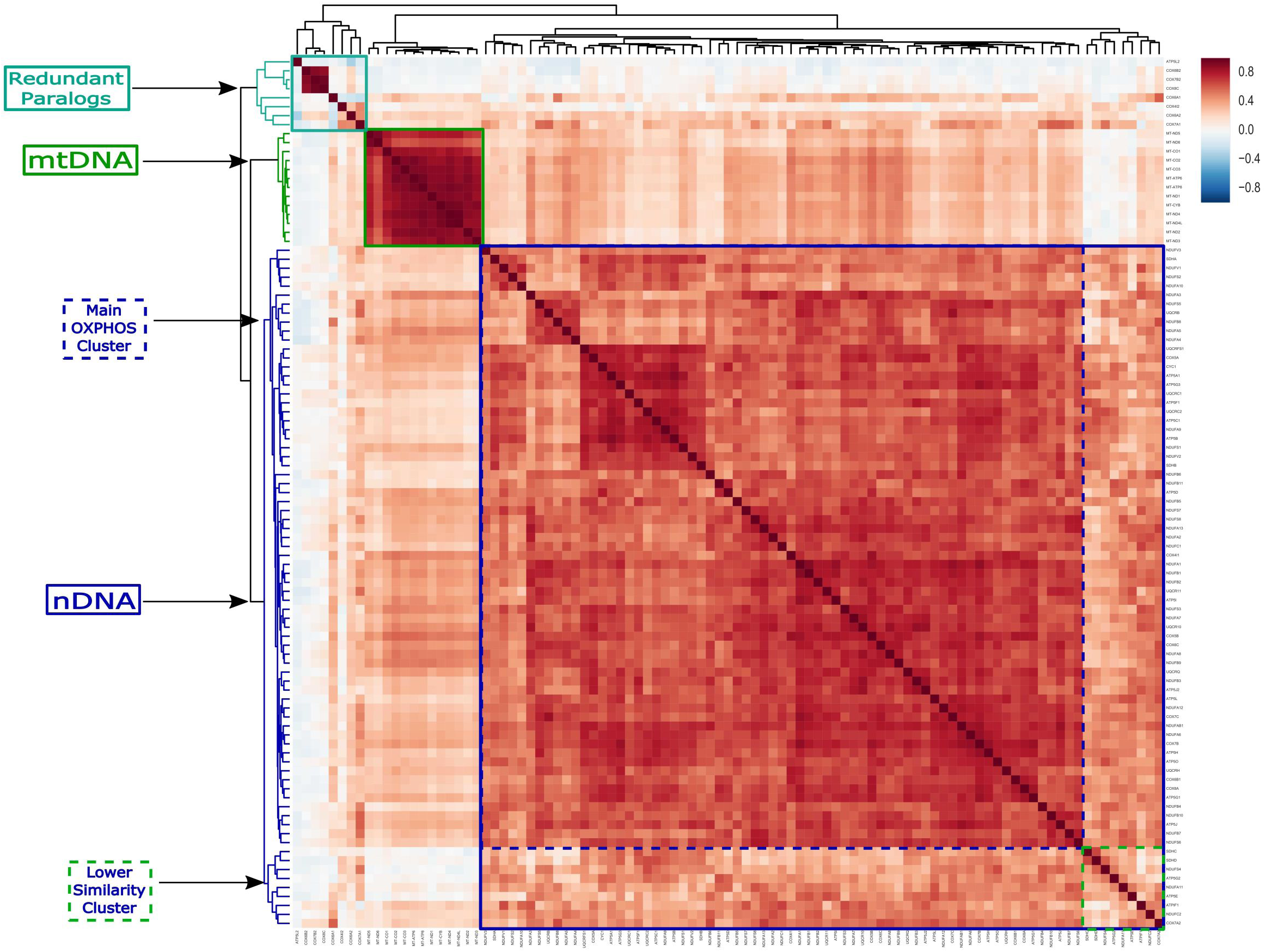
OXPHOS gene expression patterns across all tissues support the general co-regulation of mito-nuclear gene expression. A heat map of correlation values for the expression of all OXPHOS genes. The three major clusters (nDNA, mtDNA and redundant paralogs) are framed in continuous squares drawn with full lines (blue, green and cyan, respectively). More specific co-expression clusters of nDNA-encoded OXPHOS genes (main OXPHOS cluster and lower-similarity cluster) are framed with dashed lines (blue and green, respectively). A key for the color code of the Spearman’s correlation coefficients is shown in top right corner. See also Supplementary Figs. 2 and 3 and Supplementary Tables 1 and 2.

Although overall clustering of gene expression was observed between nDNA-encoded OXPHOS genes, we noticed that nine genes (*NDUFS4, NDUFC2, NDUFA11, SDHC, SDHD, COX7A2, ATP5G2, ATP5IF1* and *ATP5E*) showed relatively lower similarity to the rest of the nDNA-encoded OXPHOS genes than did others (hereafter called the ‘lower similarity cluster’) (Fig. 1). This suggested that a subset of the OXPHOS structural subunits are differentially expressed in certain tissues. To test this hypothesis, we further analyzed similarity in OXPHOS gene expression patterns in each of the 48 different tissues from the tested human subjects (see ‘Analysis of tissue-specific co-expression’ in Methods). Firstly, five of the above-mentioned genes (*COX7A2, NDUFS4, NDUFC2, NDUFA11* and *ATPIF1*) showed high similarity to the 68 nDNA-encoded OXPHOS genes in the main OXPHOS cluster (Fig. 1) only in brain tissues, consistent with some degree of tissue specificity among integral subunits of the OXPHOS system. Secondly, the expression of *COX7A1* and *COX6A2*, previously identified as heart- and skeletal muscle-specific paralogs of *COX7A2* and *COX6A1*, respectively ^25^, showed high similarity to the main OXPHOS gene cluster in 19 of the 48 tested tissues (Supplementary Fig. 2). This shared tissue-dependent co-expression pattern of *COX7A1* and *COX6A2* with the main OXPHOS gene cluster, along with the structural roles of COX7A1 and COX6A2 in complex IV dimer assembly ^26^, suggests that these subunits function as part of the OXPHOS system in a wider range of tissues than previously thought. Taken together, although general co-expression of nDNA-encoded OXPHOS genes is observed, certain nDNA-encoded OXPHOS genes vary in tissue range in which they display co-expression with the main OXPHOS gene cluster. Notably, tissue specific expression of OXPHOS transcripts is further discussed below.

### Genes coding known factors of OXPHOS protein complex assembly are not co-expressed with OXPHOS genes

We next assessed whether the high similarity between gene expression patterns of nDNA-encoded OXPHOS protein subunits is shared by known assembly factors of OXPHOS protein complexes (Supplementary Table 1). To this end, we employed the expression correlation analysis described above, this time including genes encoding assembly factors of OXPHOS complexes (Fig. 2A). In general, the correlation of tissue expression patterns between nDNA-encoded OXPHOS genes was significantly stronger than was correlation with the assembly factors (median intra-OXPHOS correlation=0.570, median OXPHOS-assembly factor correlation=0.261, MWU=8.9E4, p<1E-100) (Fig. 2B), with one exception, namely the expression pattern of assembly factor *UQCC2* clustered with 58 of the nDNA-encoded OXPHOS genes (Fig. 2A). The remaining 27 nDNA-encoded OXPHOS genes, including the eight redundant paralogs and the ‘lower similarity cluster’ genes, equally clustered with both OXPHOS and assembly factor genes. Hence, taken together, gene co-expression is highly specific for OXPHOS structural subunits (Fig. 1).

**Figure 2:**
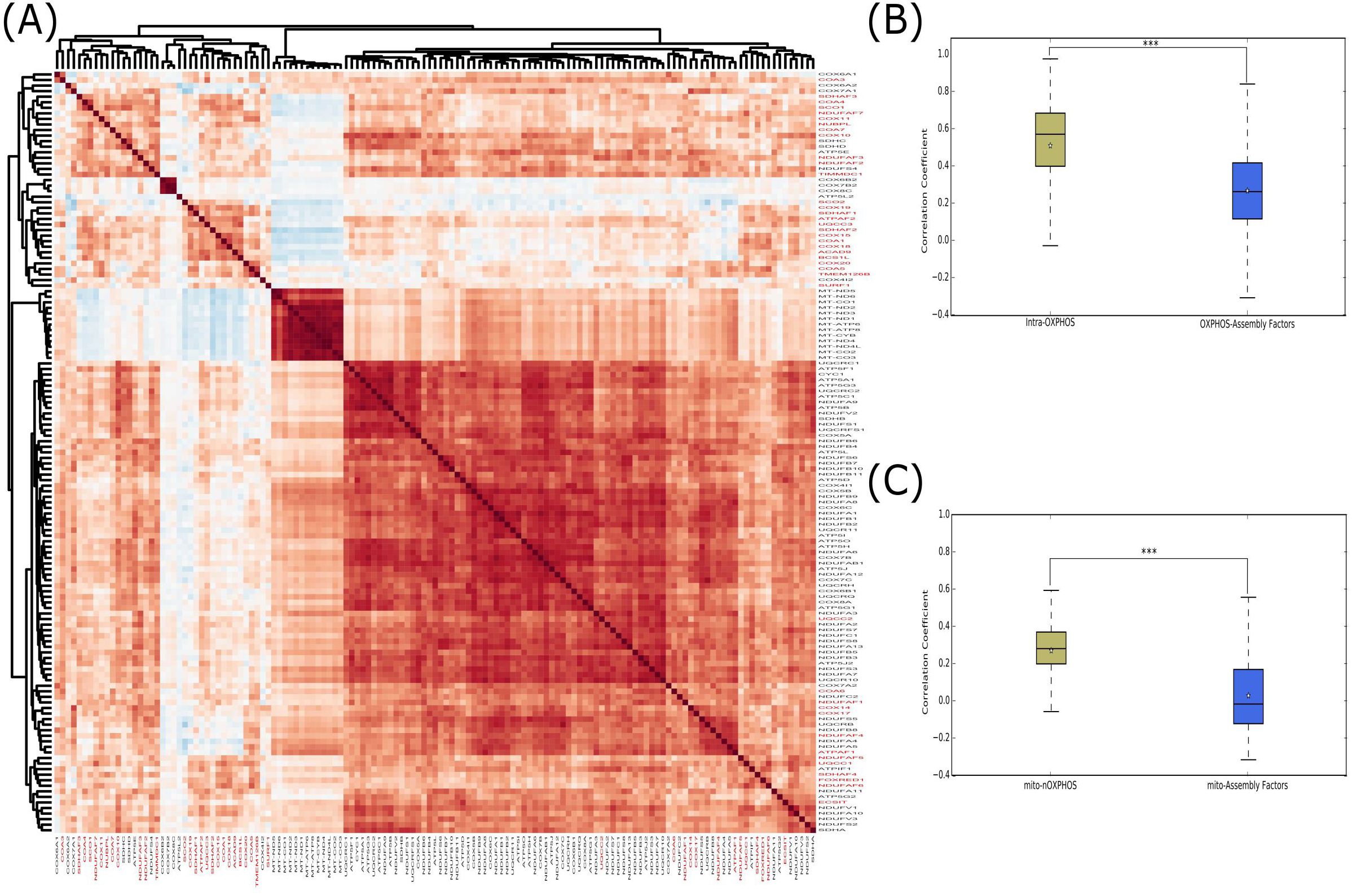
The overall positive co-expression of mitochondrial and nuclear genes is limited to OXPHOS structural subunits, excluding assembly factors. A heat map representing the Spearman’s correlations of OXPHOS gene and assembly factor expression patterns across tissues (assembly factor gene names are written in red). Genes are clustered according to similarities in their expression correlation with other genes (A). Box plots representing the correlations (Spearman’s correlation coefficients) between nDNA-encoded OXPHOS genes (intra-OXPHOS), versus their correlations with OXPHOS complexes assembly factors genes (B), and the correlations of mtDNA OXPHOS genes with nDNA OXPHOS genes, versus their correlations with OXPHOS assembly factors (C). Red lines - median correlation coefficient; stars - average correlation coefficient. *** p <1E-100. See also Supplementary Figs. 1 and 2.

### nDNA and mtDNA-encoded OXPHOS genes have correlated expression patterns

In order to analyze mtDNA gene expression while using whole transcriptomes, one has to consider that during the course of evolution mtDNA fragments migrated to the nucleus and integrated in the nuclear genome ^27,28^. Hence, it is possible that some of the RNA-seq reads may have originated from these nuclear mitochondrial pseudogenes (NUMTs). The GTEx consortium has already taken measures to identify NUMT reads and mark them as mitochondrial pseudogenes; these were excluded from further analyses. Our analysis (after NUMT exclusion) revealed high correlation between the expression patterns of all OXPHOS genes encoded by the mtDNA, with lower expression similarity observed for the *ND6* gene (Fig. 1). Such a pattern could be attributed to the polycistronic nature of mtDNA transcription: Whereas 12 out of the 13 mtDNA protein-coding genes are encoded by the heavy strand, *ND6* is the only protein-coding gene encoded by the light strand in human mtDNA.

Although forming a distinct expression cluster, the expression of all mtDNA-encoded genes was positively correlated with nDNA-encoded OXPHOS genes, apart from four redundant paralogs of OXPHOS genes (*ATP5L2, COX6B2, COX7B2* and *COX8C*) and six OXPHOS genes belonging to the ‘lower similarity cluster’ (*SDHC, SDHD, NDUFS4, ATP5G, NDUFA11* and *ATP5E*) (Fig. 1). This supports a general cross-tissue mtDNA-nDNA co-expression.

It is logical that genes encoding subunits of certain OXPHOS complexes tend to co-regulate. Indeed, in agreement with a previous report ^11^, we found that subunits belonging to the same OXPHOS complex were significantly more positively correlated than were OXPHOS genes encoding subunits belonging to different complexes (median within-complex correlation=0.625, median between-complex correlation=0.605, MWU=3.7E6, p<10E-6). Notably, such strong within-complex expression correlation was evident in OXPHOS complexes I, II, IV and V, although only marginally significant differences in correlation values were obtained for OXPHOS complex III (Supplementary Table 3).

We next compared the correlation between the expression patterns of pairs of OXPHOS genes to the correlation of OXPHOS gene expression with randomly chosen protein-coding genes. As above, each measurement was re-calculated in 1000 replicates. Our results indicate that the correlation value between OXPHOS gene pairs was significantly higher than between OXPHOS and random genes (median=0.54 and median=-0.017, respectively, MWU=0.0, p<1E-100). Notably, none of the 1000 iterations revealed a higher median correlation value between OXPHOS genes and other random protein-coding genes, as compared to the median correlation between pairs of OXPHOS genes (Supplementary Fig. 3). Similar to the case of the entire set of OXPHOS genes, the correlation of the mtDNA-encoded genes expression with nDNA-encoded OXPHOS genes (median=0.280) was significantly higher than their correlation with genes coding for OXPHOS complex assembly factors (median=-0.018, MWU=4.9E6, p<1E-100) (Fig. 2C). These findings suggest that when considering all analyzed tissues, gene expression co-regulation of mtDNA- and nDNA-encoded OXPHOS genes exists, despite the distinct mtDNA transcription regulatory system.

### mtDNA and nDNA gene expression positively correlate in all body sites apart from the ‘ancient’ brain

Although we observed positive correlation between the expression patterns of mtDNA-and nDNA-encoded genes, the medial expression of mtDNA- and nDNA-encoded OXPHOS genes in different tissues is bi-modal (Supplementary Fig. 4). We thus asked whether the general positive correlation between mtDNA- and nDNA-encoded OXPHOS genes is sustained across all tested body sites (N=48). To address this question, we analyzed each body site separately. Such analysis revealed positive correlation for all body sites, excluding several brain regions. Specifically, similar to non-brain body sites, positive mtDNA-nDNA expression correlation was observed in the neocortex (frontal cortex, cortex and cerebellar hemisphere) and the cerebellum. These regions are part of the so-called ‘mammalian’ brain ^29^. However, in most regions associated with the ‘reptilian’ brain ^30^ [hereby termed ‘ancient’ brain], which includes the hypothalamus, basal ganglia (nucleus accumbens, putamen and caudate), amygdala and the upper spinal cord (cervical c-1), we found striking negative correlation between the expression of nDNA and mtDNA-encoded OXPHOS genes (Fig. 3A). Such anti-correlation between the expression of mtDNA- and nDNA-encoded OXPHOS genes within ‘ancient’ brain regions (median=-0.563) was significantly more negative than the observed correlation between the expression of mtDNA genes and non-structural OXPHOS genes in these tissues (median=-0.098, MWU=1.7E7, p-value<1E-100) (Fig. 3B). This further suggests an ‘ancient’ brain-specific regulatory modulation, which likely impairs the positive co-regulation of OXPHOS genes encoded by the two genomes. We, however, observed several exceptions to this rule. Firstly, the substania nigra and anterior cingulate cortex (Brodmann area 24) presented a mixed/intermediate correlation pattern between the nDNA and mtDNA genes: whereas the expression of some nDNA OXPHOS genes was positively correlated with that of the mtDNA genes, some nDNA genes exhibited anti-correlation with mtDNA genes (Dataset S1). Secondly, the hippocampus, which is considered part of the ‘ancient’ brain, revealed a generally positive mito-nuclear expression correlation, apart from four nDNA genes (*SDHA, NDUFV1, NDUFS2* and *NDUFA10*), which showed somewhat negative correlation with mtDNA OXPHOS genes in this brain tissue. Notably, the expression patterns of these four genes was highly correlated across all tissues (including the ‘ancient’ brain) and did not show altered correlation to the mtDNA-encoded OXPHOS genes in the ‘mammalian’ brain tissues, thus showing some similarity between the ‘ancient’ brain and the hippocampus. Taken together, there is a strong tendency for impairment of mito-nuclear co-expression in ‘ancient’ brain tissues.

**Figure 3:**
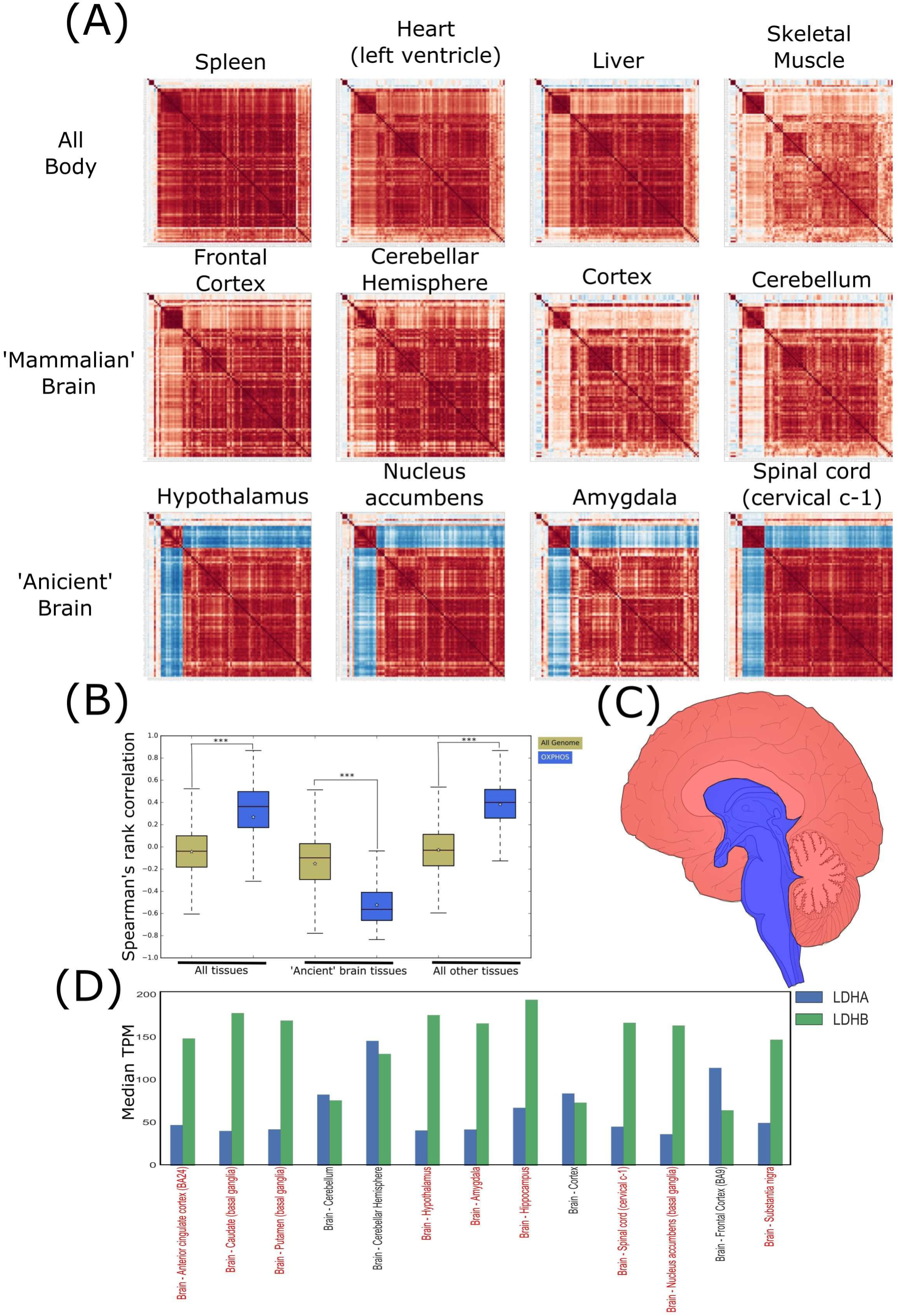
‘Ancient’ brain reveals a negative mito-nuclear expression pattern. (A) Representative tissue specific heat maps: Four representing non-brain tissues, 4 from the ‘mammalian’ brain and 4 from ‘ancient’ brain. (B) A box-plot representing Spearman’s correlation coefficients between either mtDNA genes or nDNA OXPHOS genes with the entire genome in all 48 tested tissues. The analysis is presented separately for either ‘ancient’ brain tissues or for all tissues, excluding the ‘ancient’ brain. Red lines - median correlation coefficient; stars - average correlation coefficient. *** p<1E-100. (C) A schematic illustration representing brain areas. Red zones - the ‘mammalian’ brain; Blue - ‘ancient’ brain. (D) A bar-plot showing TPM (Transcripts Per Kilobase Million) values of the two LDH subunits (LDHA and LDHB), across different human brain regions. ‘Ancient’ brain regions are indicated in red script.

### Different lactate dehydrogenase isoforms are expressed in the ‘ancient’ brain

We hypothesized that the negative correlation in gene expression between mtDNA and nDNA OXPHOS genes could reflect a limited range of assembled complexes that harbor both nDNA and mtDNA-encoded subunits at the protein level (Supplementary Table 4). This, in turn, could restrain the activity range of the OXPHOS system. One prediction that stems from this hypothesis is that ‘ancient’ brain tissues should have an altered balance between aerobic and anaerobic metabolism, relative to ‘mammalian’ brain. Such an alteration may be reflected in the expression of lactate dehydrogenase (LDH), a key enzyme that plays pivotal role in determining the fate of pyruvate. Specifically, LDH activity is important for the decision of whether pyruvate enters the tricarboxylic acid (TCA) cycle (in the form of either acetyl-CoA or oxaloacetate), thus providing electron carriers for the OXPHOS, or alternatively, whether pyruvate is fermented into lactate to maintain the NAD^+^ pool and sustain glycolysis. LDH acts as a tetramer that can contain a combination of different isoforms, including LDH-1 (the M_4_ isoform), which harbors subunits encoded only by the gene *LDHA* and is more efficient in catalyzing pyruvate reduction into lactate, and the LDH-5 (H_4_ isoform) that harbors subunits encoded only by the *LDHB* gene, which is more efficient in catalyzing lactate oxidation into pyruvate ^31,32^. Our analysis indicates that while in the vast majority of body sites (all non-brain and ‘mammalian’ brain tissues), *LDHA* and *LDHB* have similar expression levels (Supplementary Fig. 5), the ‘ancient’ brain exhibits with an increased level of *LDHA* expression (∼1.5-2.0-fold as compared to the tissue expression average) and a decrease in *LDHB* expression, to a similar extent. Interestingly, the latter pattern was also observed in the substantia nigra, the anterior cingulate cortex and the hippocampus, which have all showed some degree of anti-correlation between mtDNA- and nDNA-encoded OXPHOS genes that was not observed in the ‘mammalian’ brain tissues (Fig. 3D). These results are consistent with our suggestion, that the ‘ancient’ brain-specific mito-nuclear expression pattern correlates with a change in the expression pattern of LDH subunit genes.

### Candidate factors that best explain the co-regulation of nDNA-encoded OXPHOS genes

The identification of co-expression patterns in OXPHOS genes suggests a common underlying mechanism. Previous *in sillico* motif searches of genomic regions upstream to OXPHOS genes revealed enrichment of several transcription factor binding motifs ^11^. Although pioneering, such analysis lacked *in vivo* validation. Hence, as the first step to identifying candidates contributing to the common mechanism underlying the co-expression of OXPHOS genes, we aimed at identifying cis-regulatory elements upstream of all nDNA-encoded OXPHOS genes. To this end, we analyzed DNase-seq experimental results from 125 human cell lines generated by the Encyclopedia of DNA Elements (ENCODE) consortium and identified DNase genomic footprints (DGFs) in a window of 1000 base pairs upstream to the translation start site (TSS) of all nDNA-encoded OXPHOS genes (Supplementary Fig. 6). Then, to correlate *in vivo* binding sites of transcription factors (TFs) to the identified DGF sites, we analyzed available chromatin immunoprecipitation (ChIP)-seq results for 161 TFs from ENCODE, and assessed the overlap between the above-mentioned DGF sites and ChIP-seq peaks (Supplementary Tables 5,6 and Supplementary Fig. 6). This allowed us to identify TFs that commonly bind sites upstream to OXPHOS genes, thus providing the best candidates for explaining, at least in part, the co-expression of OXPHOS genes. Our results indicate that OXPHOS genes whose expression pattern strongly correlated also presented higher degrees of shared TF binding sites than they did with the redundant paralogs of OXPHOS genes (Fig. 1, Supplementary Fig. 6). Although our results supported some of the predictions made by van Waveren and Moraes, there were quite a few disparities. For example, van Waveren and Moraes suggested that only a low percentage (23%) of OXPHOS gene promoters harbored binding motifs for the TATA-box binding protein (TBP), and further suggested that there should be a requirement for binding of additional factors to promote transcription in TATA-less promoters. Although we identified relatively high occupancy of DGF sites upstream to many OXPHOS genes by Specificity Protein 1 (SP1) and Ying-Yang 1 (YY1) TFs (68% and 86%, respectively), associated with transcription of TATA-less promoters, we also found strong binding signals for TBP, and for the TATA Box Binding Protein-Associated Factor 1 (TAF1) in DGF sites upstream of the vast majority of OXPHOS genes (91% and 89%, respectively). Hence, in contrast to previous *in sillico* analysis, we find that TATA-binding proteins likely participate in regulating OXPHOS genes transcription.

To test for the impact of TF binding on the correlation coefficient between pairs of *in vivo*-bound nDNA OXPHOS genes (ChIP-seq), we used a least square linear regression model. Such analysis enabled the identification of 15 TFs, which bound candidate cis-regulatory elements of OXPHOS genes, and revealed a significant positive impact on the Spearman’s rank correlation coefficient between the expression patterns of OXPHOS gene pairs (Supplementary Table 7). Firstly, it is not surprising that this list included POLR2A, which *in vivo* bound 93% of the OXPHOS gene promoters and had very high impact on the OXPHOS expression correlation coefficient (P) (0.57 Spearman correlation units [SCU]). Secondly, we found *in vivo* binding signals for the histone lysine demethylase, PHD finger protein 8 (PHF8), and MYC-Associated Zinc Finger Protein (MAZ) in 86% and 81% of the OXPHOS gene promoters, respectively. Such abundant binding signals were accompanied by relatively high impact on the OXPHOS expression correlation coefficient (p) which was positively affected (0.17 and 0.056 SCU, respectively) (Fig. 4E). Thirdly, three AP-1 transcription factors had significantly positive impact on OXPHOS gene expression correlations. Specifically, JunD, FOS and FOSL2 bound 73%, 36% and 25% of the candidate OXPHOS gene promoters, respectively. These TFs also significantly affected OXPHOS gene expression pattern correlations (0.042, 0.036 and 0.048 SCU, respectively). Taken together, our analysis identified, for the first time, upstream cis-regulatory elements in all nDNA-encoded OXPHOS genes, and underlined the best candidates for mediating their co-expression.

**Figure 4:**
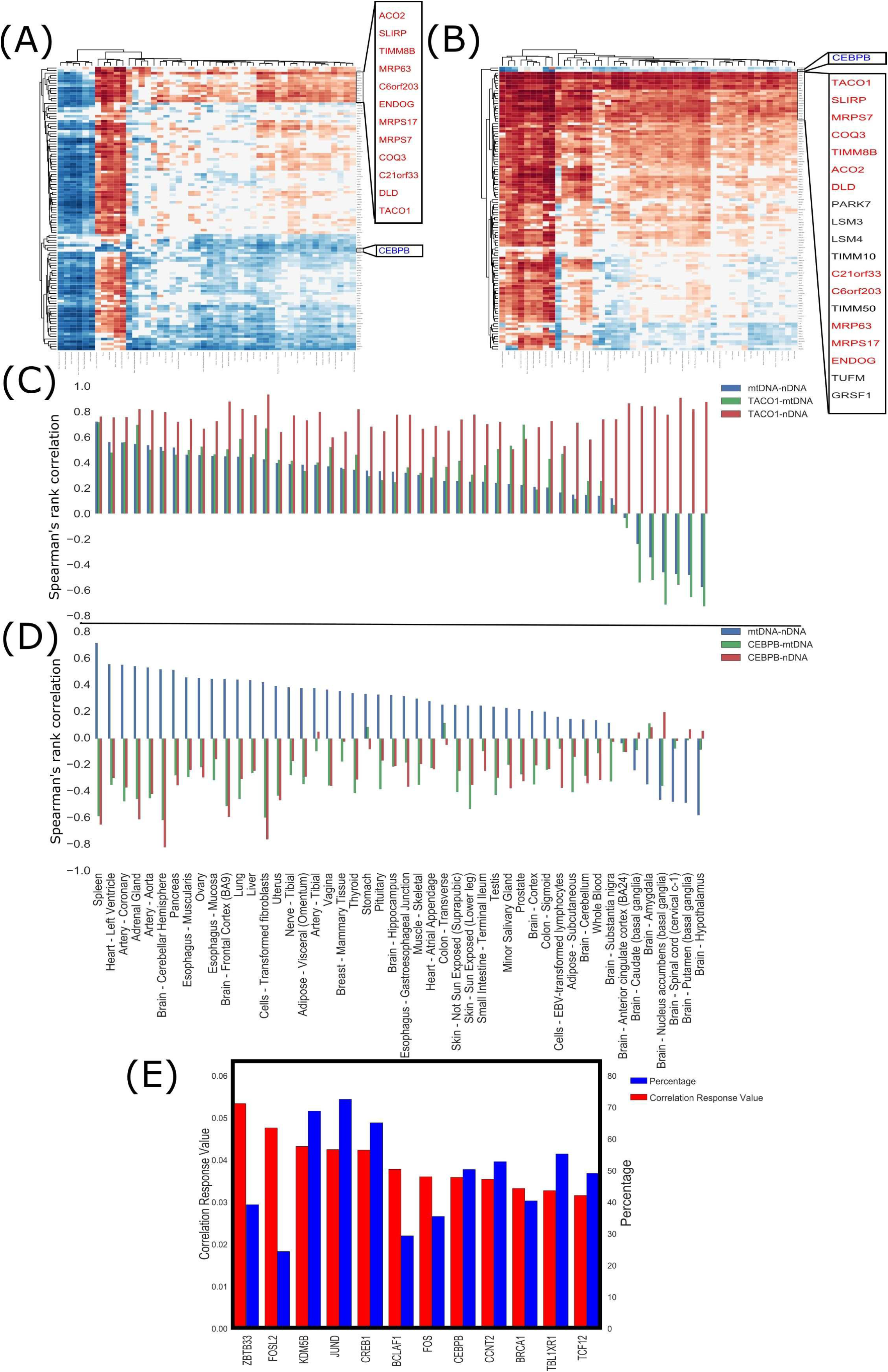
Candidate factors that best explain mito-nuclear anti-correlation in the ‘ancient’ brain. Heat maps representing the expression correlation of mitochondrial RNA-binding genes with (A) nDNA- and (B) mtDNA-encoded OXPHOS genes. Bar-plots of the expression correlations between either the mtRNA-binding gene *TACO1* (C) or the mtDNA-binding transcription factor CEBPb (D) with the median expression values of mtDNA-(green bars) and nDNA-encoded (red bars) OXPHOS genes. Blue bars represent the median value of correlation between mtDNA- and nDNA-encoded OXPHOS genes. (E) Bar-plot showing 15 transcription factors that bind cis-regulatory elements of nDNA-encoded OXPHOS genes *in vivo* and correlate with their expression. See also Supplementary Fig. 6 and Supplementary Tables 5, 6 and 7.

### Twelve RNA-binding proteins and *CEBPb* are the best candidates to explain the co-expression pattern of mtDNA- and nDNA-encoded OXPHOS genes across human body sites

We next sought for the best candidate factors to explain the co-expression of mtDNA and nDNA OXPHOS genes. To this end, we focused our analysis on a set of genes encoding both TFs and genes encoding predicted RNA-binding proteins with known mitochondrial localization in human cells. Notably, the TFs included in this list were experimentally shown *in vivo* to bind the mtDNA ^5–8,33^. To identify the factors that best explain the increase in mito-nuclear gene expression correlation, we sought factors whose expression positively correlated with OXPHOS genes encoded by both the mtDNA and the nDNA in most tissues, except for ‘ancient’ brain body sites (in which mito-nuclear correlation was negative). Twelve genes which encode predicted RNA-binding proteins (*ACO2, SLIRP, TIMM8B, MRP63, c6orf203, ENDOG, MRPS17, MRPS7, COQ3, c21orf33, DLD* and *TACO1*), showed expression correlation with both nDNA and mtDNA-encoded OXPHOS genes. Although the expression of these genes positively correlated with OXPHOS gene expression from both genomes in most tissues, their expression negatively correlated with the mtDNA genes in the ‘ancient’ brain while maintaining positive correlation with nDNA-encoded genes (Fig. 4). Interestingly, one of these genes (*SLIRP*) was previously shown to promote the stability of mRNA transcripts of both nDNA- and mtDNA-encoded OXPHOS genes ^34^. We, therefore, suggest that products of these genes serve as the best candidates to participate in post-transcriptional regulation of nDNA OXPHOS gene expression. Their negative correlation with mtDNA-encoded OXPHOS genes in ‘ancient’ brain tissues, as opposed to their positive correlation with mtDNA gene expression in other tissues, support the putative participation of these genes in active uncoupling of mito-nuclear co-regulation in the ‘ancient’ brain (Fig. 5).

**Figure 5:**
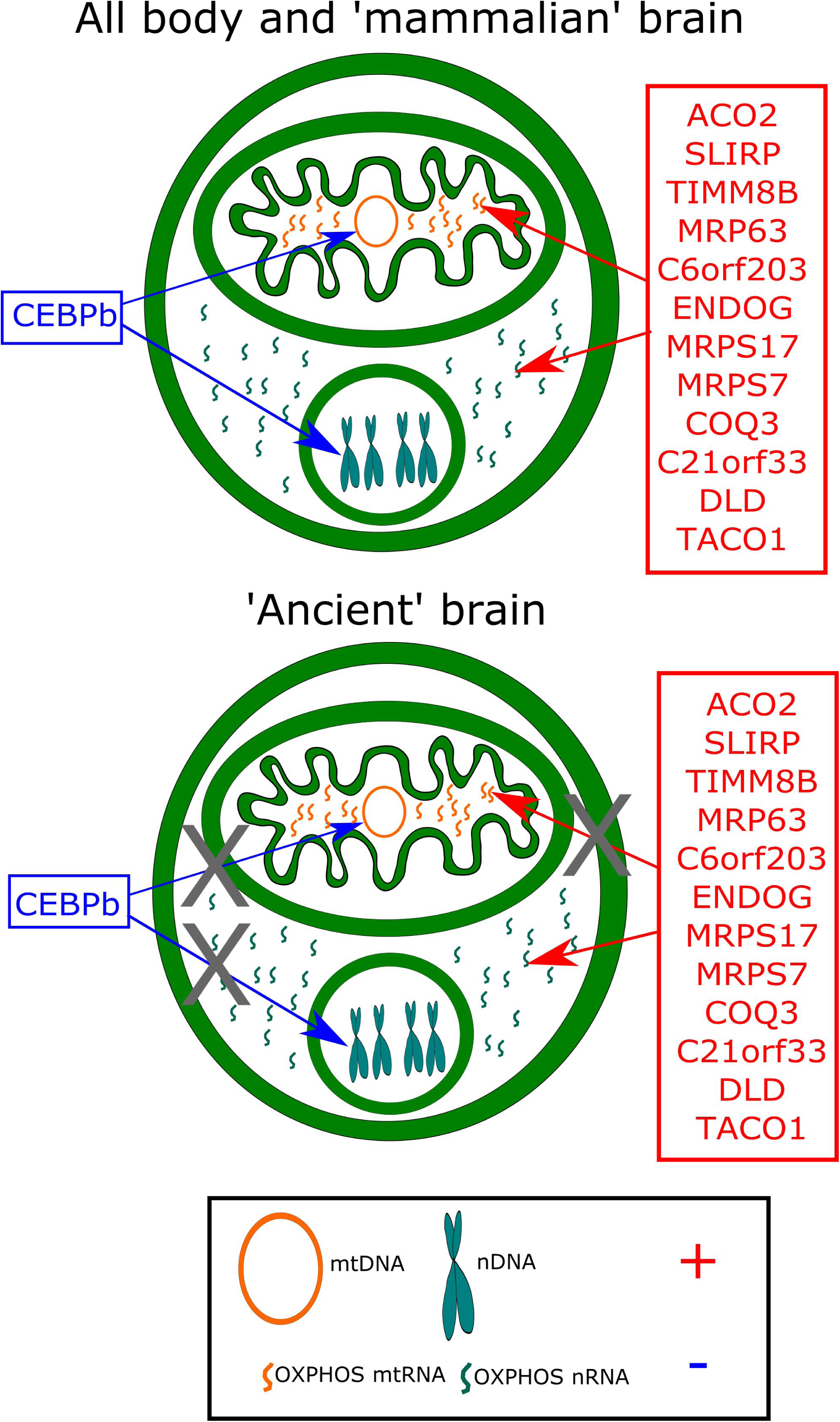
A model demonstrating the co-regulation of mtDNA and nDNA OXPHOS genes. Mitochondrial-localized RNA-binding proteins are suggested to modulate stability of mRNA OXPHOS transcripts encoded by both mtDNA and nDNA. Their anti-correlation with mtDNA-encoded genes in ‘ancient’ brain tissues provides partial explanation for the expression anti-correlation of mtDNA- and nDNA-encoded OXPHOS genes. *CEBPB* expression is negatively correlated with the expression of both mtDNA- and nDNA-encoded OXPHOS genes across human tissues but not in ‘ancient’ brain. CEBPb positively affects the expression correlation of nDNA-encoded OXPHOS genes and was shown to bind mtDNA.

Intriguingly, the expression pattern of a single gene in our list of 117 genes, *CEBPb*, was negatively correlated to the expression of both mtDNA and nDNA OXPHOS genes in most human tissues. This constitutively negative expression correlation with genes of both genomes tended to be stronger in body sites with highly positive mito-nuclear expression correlation, such as spleen, adrenal gland and the cerebellar hemisphere, but was lost specifically in the ‘ancient’ brain (Fig. 4). *CEBPb* encodes for the CCAAT/Enhancer-Binding Protein Beta (CEBPb) and was experimentally shown to modulate gene expression regulation of nDNA-encoded OXPHOS genes ^35^. We and others have shown *in vivo* binding of this protein to human mtDNA ^5,36^. We found that CEBPb *in vivo* bound 51% of the candidate promoter regions upstream to nDNA OXPHOS genes, and had a significantly positive effect on their correlation of expression patterns across human tissues (0.036 SCU). We, therefore, argue that CEBPb is likely a negative regulator (candidate repressor) of OXPHOS gene expression that concomitantly acts on both genomes to coordinate OXPHOS gene expression (Fig. 5).

### The landscape of tissue-specific expression patterns of splice variants of OXPHOS subunits

Until this point, we demonstrated co-expression of OXPHOS genes encoded by the nucleus and by the mtDNA across human tissues. Whereas most OXPHOS genes tend to co-express, we found unexpected evidence for seven OXPHOS subunits whose expression correlated with the rest of the OXPHOS genes only in a subset of tissues (Supplementary Fig. 2). Although this reduction in correlation to the main OXPHOS cluster (Fig. 1) was not tissue-specific, as compared to the known tissue-specific redundant paralogs of OXPHOS complex IV, it implied that there might be other additional attributes of tissue specificity in the expression of OXPHOS genes. Recent findings revealed tissue-dependent inclusion of two splice isoforms of NDUFV3, a structural subunit of OXPHOS complex I ^18–20^. Although interesting, it is yet unclear whether this finding reflects an exception or rather is indicative of a much wider phenomenon reflecting tissue-specific expression of splice isoforms of OXPHOS structural subunits. As a first step to assess the extent of this phenomenon, we analyzed the expression correlation patterns of *in vivo*-identified splice variants of all nDNA-encoded OXPHOS genes across and within the 48 body sites analyzed in the current study.

We analyzed expression pattern correlations among OXPHOS transcripts (including all splice variants) across all tested body sites, while employing the same analysis stages as used for analysis of gene expression patterns. Such analysis underlined the main cluster of OXPHOS gene transcripts (Fig. 6A). We found that this cluster harbors at least one splice variant representing the known canonical proteins of 63 of the 84 total nDNA-encoded OXPHOS genes (UniProt definition) (termed ‘canonical transcripts’). This ‘main OXPHOS cluster’ was used as a reference for identifying tissue-specific expression patterns of OXPHOS transcripts in a tissue-by-tissue analysis (Fig. 6B and Supplementary Dataset 2). This approach enabled the identification of canonical transcripts representing five additional OXPHOS subunits (*COX6A2, NDUFA4, NDUFC1, COX7A1* and *COX7C*) that co-expressed along with the main OXPHOS cluster in nearly a third (15) of the tested body sites. This implies that transcripts from 16 OXPHOS subunits (including the eight redundant paralogs) co-express with the main OXPHOS cluster only in certain tissues. Hence, eight non-redundant OXPHOS subunits (Fig. 1) have some degree of tissue-specific expression.

**Figure 6:**
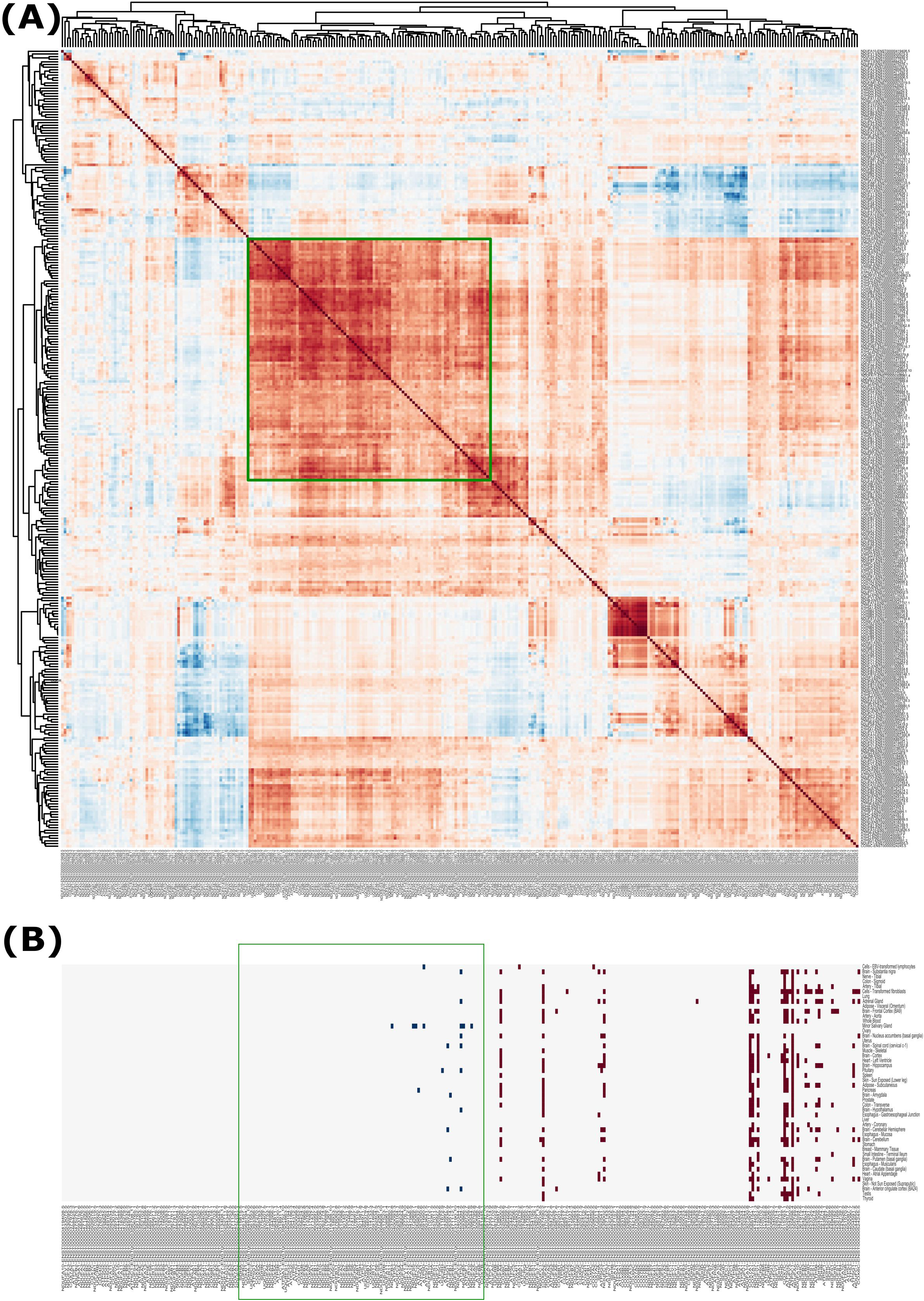
Revisiting the compendium of OXPHOS canonical subunit transcripts and identification of all tissue-specific alternatively spliced variants. (A) Heat map representing Spearman’s correlations between pairs of splice variants of nDNA-encoded OXPHOS genes. The main OXPHOS cluster is indicated by a green square. (B) A binomial heat map demonstrating similarity in the expression pattern of splice variants which are not included in the ‘main OXPHOS cluster’ to the expression of genes within this cluster in specific tissues (red). Accordingly, dissimilarities of splice variants from the ‘main OXPHOS cluster’ are indicated in blue.

Next, we closely inspected the main OXPHOS cluster, and noticed that many of the subunits were represented, either uniquely or additionally, by transcripts other than the canonical transcript. This led us to revisit the canonical sequences of all OXPHOS subunits and assess their expression patterns. To this end, we used all the UniProt canonical isoforms as query sequences to identify their co-expression with the main OXPHOS cluster in a minimum of one tissue. In the case of subunits whose known canonical isoform did not co-express with the main OXPHOS cluster in any tissue, we screened for OXPHOS co-expression of non-canonical splice isoforms representing these genes. Furthermore, in cases where the subunit sequence is known to harbor a mitochondrial localization signal, we used TargetP 1.1 to test whether the splice isoforms maintained their propensity for mitochondrial localization by more than 5% (<u>http://www.cbs.dtu.dk/services/TargetP/</u>). Our results indicate that six OXPHOS genes had such non-canonical alternatively spliced isoforms (*NDUFC2* (two alternative isoforms), *ATP5J2* (two alternative isoforms), *NDUFA2, NDUFB11, NDUFB6* and *NDUFA3*). To further consider such non-canonical transcripts as the main transcripts of these subunits, it is logical to expect that these new sequences will not affect domains of interaction with neighboring subunits in the 3D structure of the relevant OXPHOS complex. To this end, we analyzed five of the six genes with such transcripts, encoding subunits of OXPHOS complex I, for which high-resolution 3D structures from mammal sources have been recently provided ^37,38^. NDUFC2 contains two trans-membrane alpha-helixes that participate in physical interaction with the subunits ND2, NDUFC1 and NDUFA11. None of the two splice isoforms altered these two helixes. The splice isoform of *NDUFA2* is predicted to lack a single beta-strand and a single alpha-helix in the C-terminal region of the protein. However, it is unclear whether these elements play essential role in interactions with the neighboring core subunit NDUFS1. The splice variant of *NDUFB11* is not predicted to alter either of the two alpha-helixes that are likely important for interactions with ND4, NDUFB10 and NDUFB8. Neither of the two alpha-helixes of NDUFB6, including the trans-membrane alpha-helix that is important for its interaction with ND5, are altered in the alternatively spliced isoform. The two helixes in NDUFA3 that physically interact with ND1 are not predicted to change in the splice isoform either. Taken together, we propose that such non-canonical isoforms should be accepted as revised canonical isoforms of the analyzed subunits (Table 1).

## Discussion

In the current study, we quantitatively analyzed the expression of OXPHOS genes encoded by the mitochondrial and nuclear genomes in 48 human body sites. Although this detailed analysis supported correlations between OXPHOS genes, it did not support correlation of OXPHOS genes with the majority of OXPHOS complex assembly factors, suggesting that co-regulation is limited to the structural subunits of the OXPHOS system in the mitochondrial and nuclear genomes. Whereas co-expression was evident while considering all tested body sites, tissue-by-tissue analysis revealed striking disruption of the mtDNA-nDNA co-expression in tissues belonging to the ‘ancient’ brain. This finding drew our attention, especially since the more evolutionary recent parts of the brain, i.e. the cortex, cerebellar hemispheres, and cerebellum (‘mammalian brain), displayed positive mtDNA-nDNA expression correlation. This suggests that the ‘mammalian’ brain adjusted mito-nuclear co-regulation during evolution in response to its different bioenergetics requirements. Such interpretation may shed new light on the molecular basis of neurodegenerative diseases, such as Alzheimer’s disease ^39^, especially since mitochondrial gene expression alterations have been reported in various cortical tissues of Alzheimer’s disease patients.

What could be the outcome of the negative mtDNA-nDNA gene expression correlation in the ‘ancient’ brain? Analysis of mitochondrial OXPHOS activity in mouse tissues revealed a significantly higher ratio between the activity of NADH ubiquinone oxidoreductase (complex I) and the activity of succinate dehydrogenase (complex II) in the cortex than in the hypothalamus ^40^. This is interesting, since complex II contains only nDNA-encoded subunits, and complex I harbors subunits encoded by both the mitochondrial and nuclear genomes. As such, disruption of mito-nuclear co-regulation in the ‘ancient’ brain tissues may be accompanied by changes in mitochondrial activity. Secondly, mtDNA depletion in the rat striatum (an ‘ancient’ brain region) led to a significantly worse mitochondrial phenotype as compared to mtDNA depletion in the rat neocortex ^41^. These pieces of evidence suggest that in ‘ancient’ brain tissues, the amount of actively assembled mito-nuclear OXPHOS complexes (complexes I, III, IV and V) is lower than in the rest of the body (Supplementary Table 4), thus making these tissues more vulnerable to mitochondrial defects, such as reported in the striatum of Huntington’s disease patients ^42^. One can assume that given the lower range of active OXPHOS complexes in the ‘ancient’ brain, an extremely oxidative organ, this tissue requires more efficient utilization of pyruvate via the TCA cycle and, therefore, shows lower accumulation of lactate. Indeed, we found that the ratio between the expression levels of LDH subunit-coding genes (*LDHA* versus *LDHB*), was dramatically lower in ‘ancient’ brain tissues as compared to the rest of the body, including the ‘mammalian’ brain. Since LDHA favors catalyzing pyruvate-to-lactate while LDHB prefers to operate in the opposite direction (i.e., lactate-to-pyruvate), it is logical that ‘ancient’ brain tissues have a lower tendency to accumulate lactate and higher dependence on oxidative metabolism. This interpretation is supported by the observed lower tendency to accumulate lactate in ‘ancient’ brain sites (brainstem and hypothalamus), relative to the cortex, in rats after exercise ^43^. Nevertheless, proving this interpretation will require additional study.

The specific disruption of mito-nuclear co-regulation in ‘ancient’ brain tissues suggests an active regulatory phenomenon. As the first step to address this possibility, we analyzed the pattern of expression of all known regulators of mitochondrial gene expression. Such analysis revealed that the expression of certain mitochondrial RNA-binding proteins correlated with the expression of both mitochondrial and nuclear OXPHOS genes in the vast majority of human tissues while exhibiting opposite trends of correlation with mtDNA genes in the ‘ancient’ brain. It is, therefore, tempting to speculate that active un-coupling of mito-nuclear co-regulation occurred in ‘ancient’ brain tissues. This finding also shows that despite the apparently dedicated and distinct regulatory system of mtDNA gene expression, mito-nuclear co-expression was found in all non-brain tissues. This observation strongly implies the existence of regulatory cross-talk between the mitochondrial and nuclear genomes, likely through anterograde and retrograde signaling ^44^. Such signals may participate in the recently discovered translocation of nuclear transcription regulators into mammalian mitochondria, their direct mtDNA binding and their regulation of mtDNA transcription ^5,8^. Our results indicate that the expression of a gene encoding one such TF, CEBPb, is negatively correlated with the expression of both mtDNA-and nDNA-encoded OXPHOS genes in all body sites, apart from the ‘ancient’ brain, where *CEBPb* specifically lost its negative expression correlation to OXPHOS genes in both genomes. We also found that CEBPb binds regulatory elements of many nDNA-encoded OXPHOS genes and significantly affected the expression correlation between pairs of nDNA-encoded OXPHOS genes. Still, further work is required to identify the regulatory system that govern this phenomenon.

Despite the overall co-expression of OXPHOS genes, distinct clustering of mtDNA gene expression was identified in all tested tissues. This is not surprising, as core-regulators of mtDNA gene expression are dedicated to mitochondrial function and have no clear impact on nuclear gene regulation ^45^. While focusing solely on mtDNA genes, *ND6* had lower correlation scores with rest of the genes encoded by this genome. This is consistent with *ND6* being the only mRNA encoded by the light mtDNA strand, which in humans has a separate promoter from the heavy mtDNA strand, that harbors most mtDNA genes. It would, therefore, be of great interest to assess correlation in mtDNA gene expression in birds and amphibians, where a bidirectional mtDNA promoter governs the transcription of both mtDNA strands ^46,47^. As our expression correlation was observed at the population level, this analysis awaits the availability of RNA-seq from multiple unrelated individuals from these taxa.

RNA-seq analysis enabled the identification of both canonical and non-canonical splice variants of all nDNA-encoded OXPHOS genes, and assessment of their expression patterns in human body sites. Our analysis showed that while most OXPHOS genes co-expressed in most tissues, such co-expression clusters were not always represented by a combination of the known canonical gene transcripts. Close inspection revealed that six genes encoding OXPHOS subunits, including *NDUFC2, ATP5J2, NDUFA2, NDUFB11, NDUFB6* and *NDUFA3*, were mostly represented by a different, non-canonical splice variant, resulting in a non-canonical protein. Therefore, based on clustering and abundance in the different tissues, we offer a revised list of OXPHOS canonical transcripts, which will be useful for future analyses of OXPHOS genes (Table 1). Our analysis also revealed that the expression of nearly a quarter of all OXPHOS subunits was either clearly tissue-specific or correlated with the main OXPHOS co-expression cluster only in a subset of tissues. This suggests that the subunit composition of several OXPHOS complexes varies among tissues. This hypothesis is supported by the existence of tissue-specific paralogs of cytochrome C-oxidase (complex IV) ^48,49^ and by the recent identification of a tissue-specific splice variant of the complex I subunit *NDUFV3*, also at the protein level ^18–20^. Notably, such interpretation awaits proteomics analysis of multiple human tissues that potentially will differentiate paralogs from splice isoforms. Regardless, our analyses provide a first glance into the landscape of the tissue expression pattern of OXPHOS subunit splice variants, and constitutes the first assessment of this phenomenon.

**Table 1:** The revisited list of canonical OXPHOS splice variants.

| Subunit | Revised Canonical Transcript (Ensemble ID) | Revised Inferred Protein (UniProt ID) | Previous Canonical Transcript (Ensemble ID) | Previous Inferred Protein (UniProt ID) |
| --- | --- | --- | --- | --- |
| **NDUFA2** | ENST00000512088.1 | O43678-2 | ENST00000252102.8 | O43678-1 |
| **NDUFA3** | ENST00000391763.3 | A8MUI2 | ENST00000485876.5 | O95167 |
| **NDUFB6** | ENST00000350021.2 | O95139-2 | ENST00000379847.7 | O95139-1 |
| **NDUFB11** | ENST00000276062.8 | Q9NX14-2 | ENST00000377811.3 | Q9NX14-1 |
| **NDUFC2** | ENST00000528164.1 | E9PQ53-1 | ENST00000281031.4 | O95298-1 |
|  | ENST00000525085.1 | O95298-2 |  |  |
| **ATP5J2** | ENST00000394186.3 | P56134-2 | ENST00000292475.7 | P56134-1 |
|  | ENST00000359832.8 | P56134-3 |  |  |

In summary, to the best of our knowledge, we have presented here the most comprehensive analysis to date of the OXPHOS transcription regulatory landscape across multiple human tissues. We discovered overall co-regulation of the mitochondrial and nuclear genomes, which was disrupted in the ‘ancient’ brain but not in the ‘mammalian’ brain, possibly due to an uncoupling of mitochondrial post-transcriptional regulation and by a loss of transcription coordination of both genomes. We interpreted this finding as reflecting adaptation of the ‘mammalian’ brain to newly emerging energy needs during evolution. Our detailed analysis of cis-regulatory elements of all nDNA-encoded OXPHOS genes revealed several candidate transcription and post-transcription regulators that might provide a partial explanation to the observed co-regulation of OXPHOS subunits. Finally, we revisited the list of canonical OXPHOS transcripts and provided a comprehensive view of tissue-specific transcripts, thus suggesting tissue variability not only in the regulatory pattern but also in the composition of OXPHOS complexes.

## Materials and Methods

### Source of RNA-seq data in RPKM (Reads Per Kilobase per Million mapped reads)

Gene expression data and individual information, including gender and cause of death, were obtained from the publicly available data at the GTEx portal (<u>www.gtexportal.org</u>). Individual ages (rather than age groups) were obtained from the GTEx-authorized data table for donor information (pht002742.v6.p1.c1), ‘age’ column (phv00169063.v6.p1.c1).

### Extraction of alternatively spliced isoforms

The expression patterns of splice variants of the tested genes were extracted from the publically available data at the GTEx portal. Only protein-coding variants, determined according to the Ensemble genome browser (<u>www.ensembl.org</u>), were considered in our analysis of co-expression.

### NUMT exclusion

One source of contamination of our analyzed mtDNA gene expression data is sequencing reads of ancient mtDNA fragments that were transferred to the nDNA during the course of evolution ^27,28^. To control for such contamination, GTEx used nuclear mitochondrial pseudogene transcript sequences as their reference transcriptome (appears as mtXPN in the GTEx data, where X represents the mitochondrial gene and N stands for the number of the pseudogene). Therefore, reads that had higher similarity scores to any given pseudogene, as compared to the reference mtDNA gene, were excluded from the reads count for mtDNA-encoded genes. To further control for NUMTs, we used the normalized read counts (reads per kilo-base per million reads - RPKM) available in the GTEx consortium data files as our raw data. According to GTEx documentation, the read alignment distance was ≤6 (i.e., sequence alignments must not contain more than six mismatches), thus notably reducing possible contamination by NUMT reads.

### RPKM to TPM conversion

The RPKM value for a given gene “i” in each library (each sample) was converted to TPM as previously described ^50^, using the following formula:

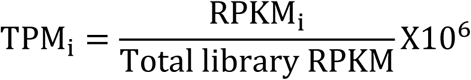

### Control for age, gender and cause of death

To control for the possible contribution of age, gender and cause of death to gene expression patterns, we normalized for these factors using linear regression (see below), such that the calculated residual TPM reflected age/gender/cause-of-death-independent gene expression. Notably, we compared the vectors of residual TPMs to obtain Spearman correlation scores between each pair of tested genes.

### Gene/transcript specific linear regression

To obtain the residual gene expression (i.e., age/gender/cause of death-independent values), we considered all these factors as independent regressing agents in a least-square linear regression model. The age variable was obtained as a continuous factor, whereas gender and cause-of-death were categorical factors. Dummy variables (1 for gender and 5 for cause-of-death) were used to correct for the effect of these factors on gene expression. Finally, the residual TPM for each gene in each sample was calculated as the difference between the observed TPM and the TPM value predicted for this gene by the linear regression, considering phenotypic parameters (i.e. age, gender and cause-of-death).

### Secondary structure predictions

To assess the possible impact of any changes of interest in structural elements that are predicted to be important for the interaction of a tested OXPHOS protein with its partners, we used Jpred 4 (<u>http://www.compbio.dundee.ac.uk/jpred/</u>).

### Identification of digital genomic footprinting sites upstream to OXPHOS genes, and their occupation by transcription factors

DNase-seq experiments performed by the ENCODE consortium were analyzed in terms of a 1000 bp region upstream to the translation start site of all genes encoding OXPHOS subunits and assembly factors in the nuclear genome. The DNase-seq foot-printing data was taken from ENCODE DNaseI Hypersensitive Site Master List (<u>http://genome.ucsc.edu/cgi-bin/hgTrackUi?db=hg19&g=wgEncodeAwgDnaseMasterSites</u>).

Translation start sites (TSS) were annotated according to the GRCh37/hg19 RefSeq genes track and were taken from the UCSC table browser (<u>https://genome.ucsc.edu/cgi-bin/hgTables</u>). Transcription factor-binding sites were obtained from the ENCODE clustered TFs list (<u>http://genome.ucsc.edu/cgi-bin/hgTrackUi?db=hg19&g=wgEncodeRegTfbsClusteredV3</u>). For each OXPHOS gene, a positive TF-binding site was listed if its binding peak was identified within the 1000 bp window upstream to the TSS and overlapped with an identified DNase-seq footprinting site/s.

### Analysis of co-expression

To assess co-expression between genes across the available multiple tissues in the GTEx database, we focused our analysis on 48 tissues, for which a minimum of 50 samples had available RNA-seq data. Then, we sampled 50 samples from each tissue and calculated Spearman’s rank correlation coefficients between pairs of genes\transcripts according to their 2400 (50 times 48) residual TPM values. We repeated this calculation a 1000 times and considered the correlation to be significant only if the P-value for the Spearman’s rank correlation was smaller than 0.05 in a minimum of 950 iterations (95% of the iterations). The median correlation coefficient of each pair of genes/transcripts, obtained after the 1000 random iterations, was considered as the median Spearman’s rank correlation coefficient for this pair (Supplementary Fig. 1).

### Clustered heat maps construction

The median Spearman’s rank correlation coefficient between the pairs of genes/transcripts under consideration were used to create a distance matrix for hierarchical clustering. Such clustering was obtained according to the similarity of the correlation of gene/transcript expression with other genes/transcripts under consideration. The correlation values were color-coded: red – positive correlation, blue – negative correlation. Non-significant correlations are in white.

To construct heat maps for each of the analyzed tissues, we used the gene/transcript order that was obtained from the hierarchal clustering generated while analyzing the total expression pattern across all tissues. Additionally, while constructing tissue-specific correlation distance matrixes, we considered all samples available for the tested tissue.

### Analysis of tissue-specific co-expression

For each gene/transcript, we calculated the expression correlation, and the median of these correlations coefficients (M), with the main OXPHOS cluster (Figs. 2 and 6) across tissues and within each tissue. We marked the number of these positive correlations by ‘N’. Expression of a gene/transcript was considered highly similar to the expression of the main OXPHOS cluster in a given tissue only if the product of the number of positive correlations with the main OXPHOS cluster and the median of these correlations coefficients (N X M) in this tissue was higher than the median N X M value of genes/transcripts of the main OXPHOS cluster across all tested tissues. Accordingly, a gene/transcript included in the main OXPHOS cluster according to our analysis was considered as having low similarity to the main OXPHOS cluster in a given tissue if its N X M value in this tissue was lower than the median N X M value of the genes/transcripts outside the main OXPHOS cluster across all tissues.

### Statistical tools and data visualization

All statistical analyses and data visualization were performed using Python 2.7 scipy.stats, pandas, matplotlib and seaborn packages (Dataset S3). Validations to the performance of the Python scripts were performed using STATISTICA 12.

## Acknowledgements

The authors express their deep appreciation to Dr. Alal Eran, Prof. Hermona Soreq, Dr. Esti Yeger-Lotem, Dr. Barak Rotblat, Prof. Ofer Ovadia, Prof. Avraham Zangen and Dr. Noam Barnea-Yigael for deep and insightful discussions on the brain-specific expression pattern found in this study. This study was funded by grants from the Binational Research Foundation and US Army Life Sciences division awarded to DM. The authors also acknowledge the Darom scholarship for excellent PhD students awarded to GB.

## Author Information

GB performed the majority of data analyses, AB analyzed the TF-binding site and DNase-seq data. DM conceived the study. DM and GB wrote the paper.

## Supplemental Information

### Supplemental Figures

**Supplementary Figure 1. Workflow of the expression pattern correlation analysis.** See also dataset S3 for Python scripts used during the analysis.

**Supplementary Figure 2. Tissue-dependent OXPHOS gene expression.** A clustered binomial heat map demonstrating similarity in the expression pattern of OXPHOS genes which are not included in the ‘main OXPHOS cluster’ to the expression of genes within this cluster in specific tissues (red). Dissimilarities of OXPHOS genes from the ‘main OXPHOS cluster’ are indicated in blue. The orange and green rectangles show the two genes groups, (COX7A1 and COX6A2) and (COX7A2, NDUFS4, NDUFC2, NDUFA11 and ATPIF1), respectively. The gray rectangles show brain tissues.

**Supplementary Figure 3. nDNA-encoded OXPHOS genes show significantly higher co-expression with mtDNA-encoded OXPHOS genes**. A box-plot showing the values of the correlation coefficients between pairs of OXPHOS genes (intra-OXPHOS) and between OXPHOS genes and randomly chosen protein-coding genes (OXPHOS-genome). *** p <1E-100.

**Supplementary Figure 4. The correlation distribution of nDNA- and mtDNA-encoded OXPHOS genes is bi-modal.** A histogram comparing the expression distribution of nDNA-encoded OXPHOS genes (green) and all other genes in the genome (blue) with medial mtDNA expression in all tissues.

**Supplementary Figure 5. Expression of LDH subunits across human tissues.** A bar-plot showing the median TPM values of the two LDH subunit-coding genes (LDHA and LDHB) across all human tissues inspected in this study.

**Supplementary Figure 6. Graphical representation of the *in vivo* identification of candidate promoters in nuclear DNA-encoded OXPHOS genes.** (A) A map of the upstream region of a representative OXPHOS subunit (NDUFA8) tested for TF binding in DNaseI Genomic Footprinting (DGF) sites. X-axis represent the positions in the promoter region, relative to the translation start site (TSS). DGF sites are indicated by blue lines in the upper row and below, all the listed TFs binding to the 1000 bp promoter region are shown, with their ChIP-seq peaks shown as black lines. Note that only TFs with a ChIP-seq peak overlapping a minimum of one DGF are indicated. (B) Representation of the results for all nDNA-encoded OXPHOS genes. X-axis represent the positions in the promoter region, relative to the translation start site (TSS). DGFs are shown by blue lines if they do not overlap with any identified TF-binding peak in the ChIP-seq experiments and in black and blue lines if they overlap with at least one such TF-binding peak.

### Supplemental Tables

**Supplementary Table 1.** A list of OXPHOS genes analyzed in this study, following HUGO Gene Nomenclature Committee (HGNC). OXPHOS structural and assembly factors are indicated, along with their Ensemble accession numbers.

**Supplementary Table 2.** Sub-division of OXPHOS structural genes into tissue expression clusters.

**Supplementary Table 3.** Summary statistics of the expression differences between subunits within or between OXPHOS complexes.

**Supplementary Table 4.** A mathematical representation of the effect of negative expression correlation between mtDNA- and nDNA-encoded OXPHOS genes on the OXPHOS complex assembly range. M_0_, N_0_ = Initial expression values of mtDNA (M_0_) and nDNA (N_0_) genes, ΔM, ΔN = Changes in expression of mtDNA (ΔM) and nDNA (ΔN) genes.

**Supplementary Table 5.** Coordinates of DNase I footprinting sites upstream to all human OXPHOS genes.

**Supplementary Table 6.** OXPHOS structural genes and the list of TFs that bind within DGFs in their candidate cis-regulatory regions.

**Supplementary Table 7.** Statistically significant Spearman’s rank correlation coefficient positively affecting TFs. * Candidate promoter regions.

### Supplemental Datasets

**Supplementary Dataset 1**: Correlation heat maps for OXPHOS gene expression patterns in 48 human tissues. <u>https://figshare.com/s/71045c9d0abe3e82866c</u>

**Supplementary Dataset 2:** Correlation heat maps for expression patterns of splice variants of OXPHOS genes in 48 human tissues. <u>https://figshare.com/s/c2342a1e655467e57ab2</u>

**Supplementary Dataset 3:** Python scripts for raw data conversion, normalization and co-expression analysis. <u>https://figshare.com/s/cd1a39a7359580a1a4db</u>

